# Paired-Sample and Pathway-Anchored MLOps Framework for Robust Transcriptomic Machine Learning in Small Cohorts

**DOI:** 10.1101/2025.06.03.657721

**Authors:** Mahdieh Shabanian, Nima Pouladi, Liam S. Wilson, Mattia A. Prosperi, Yves A. Lussier

## Abstract

**Background:** Ninety percent of the 65,000 human diseases are infrequent, collectively affecting ∼ 400 million people, substantially limiting cohort accrual. This low prevalence constrains the development of robust transcriptome-based machine learning (ML) classifiers. Standard data-driven classifiers typically require cohorts of over 100 subjects per group to achieve clinical accuracy while managing high-dimensional input (∼25,000 transcripts). These requirements are infeasible for micro-cohorts of ∼20 individuals, where overfitting becomes pervasive.

**Objective:** To overcome these constraints, we developed a classification method that integrates three enabling strategies: (i) paired-sample transcriptome dynamics, (ii) N-of-1 pathway-based analytics, and (iii) reproducible machine learning operations (MLOps) for continuous model refinement.

**Methods:** Unlike ML approaches relying on a single transcriptome per subject, within-subject paired-sample designs — such as pre-versus post-treatment or diseased versus adjacent-normal tissue — effectively control intra-individual variability under isogenic conditions and within-subject environmental exposures (e.g. smoking history, other medications, etc.), improve signal-to-noise ratios, and, when pre-processed as single-subject studies (N-of-1), can achieve statistical power comparable to that obtained in animal models. Pathway-level N-of-1 analytics further reduces each sample’s high-dimensional profile into ∼4,000 biologically interpretable features, annotated with effect sizes, dispersion, and significance. Complementary MLOps practices—automated versioning, continuous monitoring, and adaptive hyperparameter tuning—improve model reproducibility and generalization.

**Results:** In two case studies—human rhinovirus infection versus matched healthy controls (n=16 training; 3 test) and breast cancer tissues harboring TP53 or PIK3CA mutations versus adjacent normal tissue (n=27 training; 9 test)—this approach achieved 90% precision and recall on an unseen breast cancer test set and 92% precision with 90% recall in rhinovirus fivefold cross-validation. Incorporating paired-sample dynamics boosted precision by 8.8% and recall by 6%, while the MLOps workflow yielded additional gains of 14.5% and 12.5%, respectively. Moreover, our method identified 42 critical gene sets (pathways) for rhinovirus response and 21 for cancer mutation status.

**Conclusions:** These proof-of-concept results support the utility of integrating intra-subject dynamics, “biological knowledge”-based feature reduction (pathway-level feature reduction grounded in prior biological knowledge; e.g., N-of-1-pathways analytics), and reproducible MLOps workflows can overcome cohort-size limitations in infrequent disease, offering a scalable, interpretable solution for high-dimensional transcriptomic classification. Future work will extend these advances across various therapeutic and small-cohort designs.

## 1 Introduction

Precision medicine seeks to personalize healthcare by accounting for individual differences in genetic makeup, environmental exposures, and lifestyle factors. This tailored approach becomes especially challenging when analyzing high-dimensional transcriptomic data derived from small patient cohorts (micro-cohorts), a scenario frequently encountered in studies of rare or infrequent diseases. Micro-cohorts typically involve datasets characterized by high dimensionality (approximately 25,000 transcriptomic features) juxtaposed against limited sample sizes (approximately 20 subjects), conditions that commonly induce overfitting in traditional machine learning models. Advanced analytical methodologies have thus become essential in identifying robust and clinically meaningful biomarkers from these small-scale studies to facilitate personalized patient care.

A large share of the ∼65,000 known human diseases are infrequent—neither rare nor common— making it difficult to assemble statistically robust cohorts without multi-year, multi-center efforts. Around 5.9% of the global population is affected by rare diseases (EURORDIS, Organization for Rare Diseases, 1), highlighting their substantial impact on global health.

In highly heterogeneous diseases, such as cancer, where tumor subtypes and genetic mutation profiles can vary substantially between individuals, conventional machine learning approaches often suffer from insufficient statistical power and heightened risk of overfitting. To mitigate these challenges, N-of-1 analytics has emerged as an innovative approach, allowing individuals to serve effectively as their own controls. By measuring within-subject transcriptomic changes and integrating these measurements into biologically interpretable pathway-level features, N-of-1 analyses significantly reduce noise and enhance the detection of biologically meaningful signals, even amidst substantial inter-subject variability (2-8).

Concurrently, the emergence of machine learning operations (MLOps), inspired by DevOps practices, has significantly improved the deployment, optimization, and monitoring of machine learning models. MLOps leverages automated experiment tracking, hyperparameter tuning, and continuous integration, enhancing workflow efficiency, reliability, reproducibility, and scalability— factors essential for developing robust and maintainable models in biomedical research (9-15).

We hypothesized that integrating three complementary strategies would enhance classification accuracy and robustness in micro-cohort scenarios: (i) implementing MLOps frameworks to achieve robust and reproducible model performance, (ii) leveraging transcriptomic dynamics observed between paired biological samples (e.g., diseased versus healthy tissues from the same individual), and

(iii)utilizing single-subject analytics (N-of-1-pathway analysis) to generate biologically interpretable features, namely approximately 4,000 human-curated biological pathways along with their respective effect sizes and statistical significance.

To empirically test this hypothesis, we conducted a proof-of-concept analysis on two distinct human micro-cohorts, each comprising paired biological samples representing two different tissue conditions per subject. For each cohort, we systematically evaluated three distinct data transformation strategies: (1)conventional analysis using only the affected tissue per subject, (2) fold-change transformation involving the ratio of affected tissue mRNA expression to paired control tissue expression for each subject, and (3) N-of-1-pathway transformation, summarizing individual subject-level pathway effect sizes and p-values.

Each of these three data transformations was subjected to classification modeling both with and without incorporating MLOps, resulting in a total of twelve experimental conditions across both cohorts. To further validate the robustness and relevance of features selected by the best-performing classifier, we conducted a rigorous retrospective ablation analysis. Specifically, in ablation analysis, we masked the top twenty discriminative features from the dataset and assessed the resulting impact on classification accuracy and stability. This comprehensive analysis framework allowed us to quantify the individual contributions of key biomarkers to the model’s predictive performance.

## 2 Method and Material (Figure 1)

### 2.1 Human Cohort Datasets (Table 1)

Datasets (**Table 1**) were selected for their spanning cancer and infection, with small sample sizes, varying heterogeneity, and paired tissue samples per subject. Processing followed published methods, ensuring prior studies comparability (16, 17).

**Table 1.**
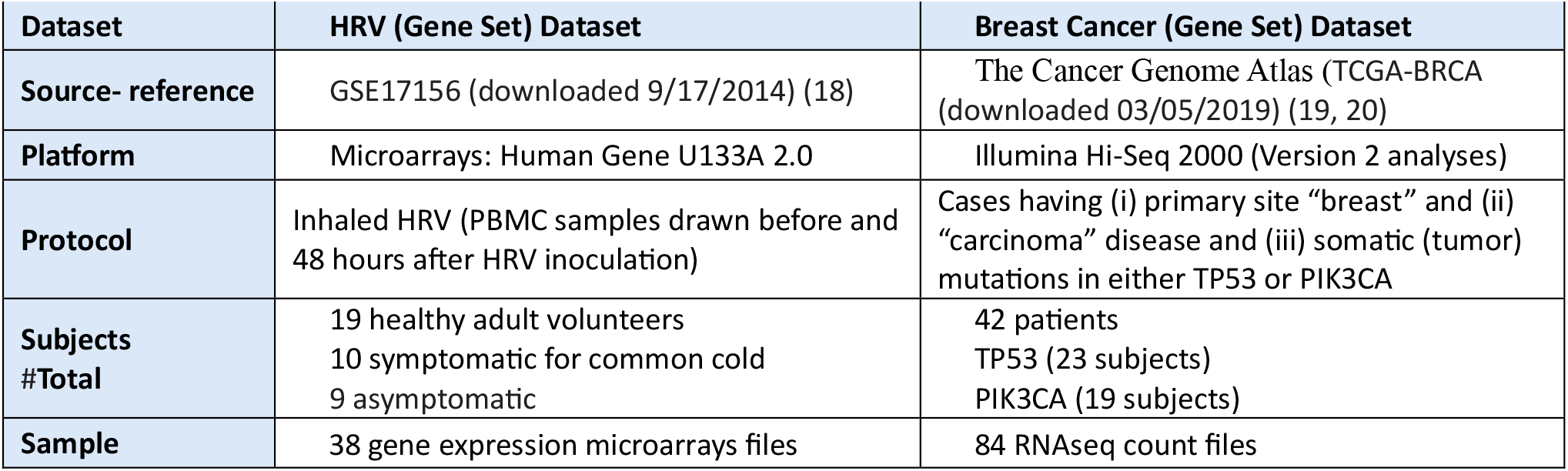
Description of the two Human Cohort Datasets. Legend: # = count of subjects;

| Dataset | HRV (Gene Set) Dataset | Breast Cancer (Gene Set) Dataset |
| --- | --- | --- |
| Source- reference | GSE17156 (downloaded 9/17/2014) (18) | The Cancer Genome Atlas (TCGA-BRCA (downloaded 03/05/2019) (19, 20) |
| Platform | Microarrays: Human Gene U133A 2.0 | Illumina Hi-Seq 2000 (Version 2 analyses) |
| Protocol | Inhaled HRV (PBMC samples drawn before and 48 hours after HRV inoculation) | Cases having (i) primary site “breast” and (ii) “carcinoma” disease and (iii) somatic (tumor) mutations in either TP53 or PIK3CA |
| Subjects #Total | 19 healthy adult volunteers<br>10 symptomatic for common cold<br>9 asymptomatic | 42 patients<br>TP53 (23 subjects)<br>PIK3CA (19 subjects) |
| Sample | 38 gene expression microarrays files | 84 RNAseq count files |

**Fig 1.**
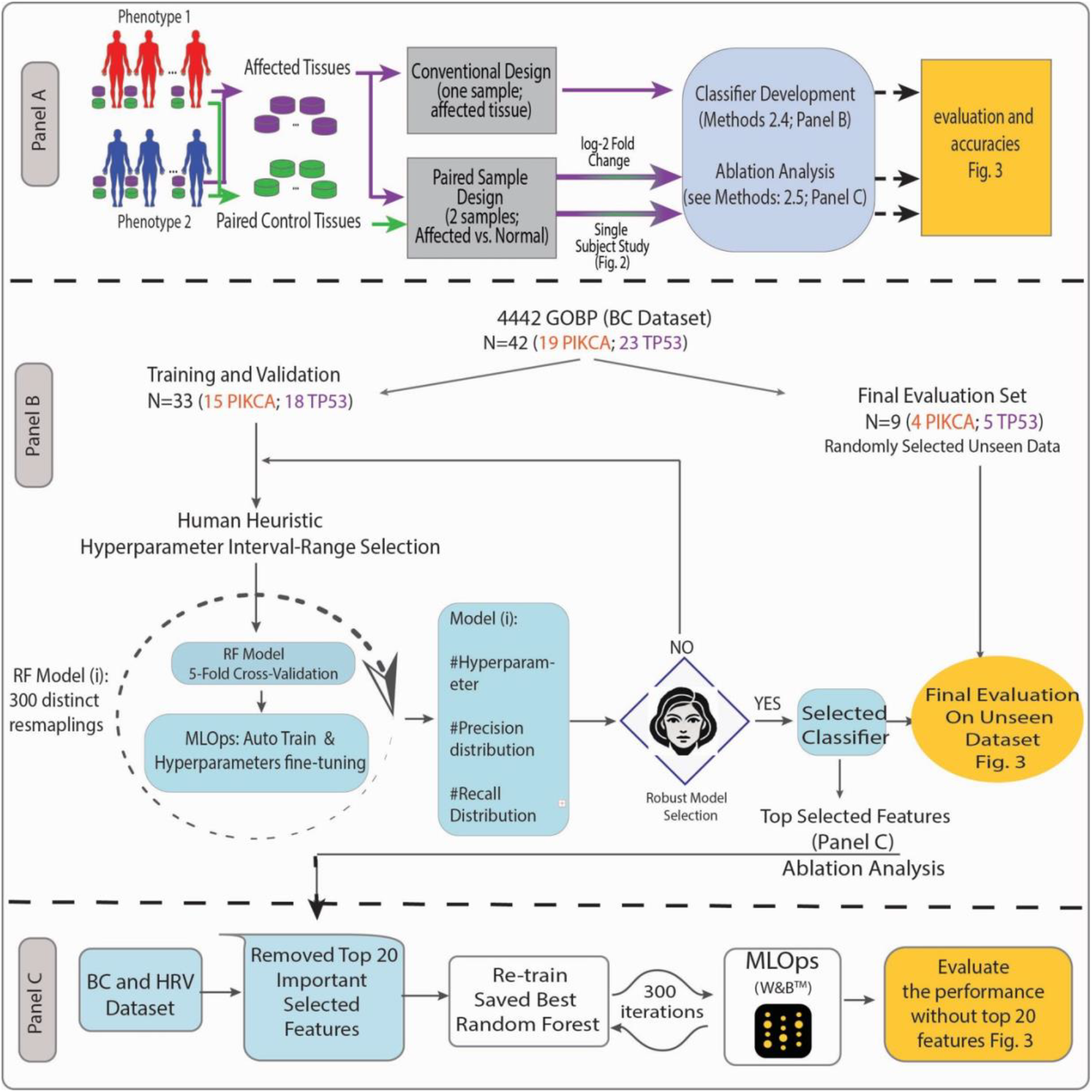
<u>Panel A</u>. Overview of Methods and Process Flow of the Proof-of-Concept Study. Classification methods are applied to two cohorts (**Table 1**), each with two distinct clinical phenotypes: (i) Breast Cancer (BC) subjects, stratified by oncogenic drivers (TP53 vs. PIK3CA), and (ii) Human Rhinovirus (HRV)-infected subjects (symptomatic vs. asymptomatic). Each subject provides two samples under different conditions: (i) BC—within-subject comparison of cancerous tissue vs. unaffected margins, and (ii) HRV—within-subject comparison before vs. during infection. Six classification experiments are conducted on each cohort’s extracted transcriptomes, evaluating three complementary classification strategies for micro-cohorts: (1) MLOps-driven robustness, (2) transcriptome dynamics between paired samples (e.g., exposed vs. unexposed tissue), and (3) single-subject pathway analytics (N-of-1; detail**s** in **Figure 2**). **<u>Panel B</u>**. Random Forest (RF) Classifier Pipeline in the BC Cohort. The RF classification workflow consists of three key steps after extracting an unseen evaluation set: (1) hyperparameter tuning using Weights & Biases MLOps, (2) iterative model refinement via 300 resampling cycles of five-fold cross-validation (80% samples in the training set, 20% in the validation set), and (3) human-in-the-loop expert review to assess failure patterns and overfitting. The expert either adjusts hyperparameters for another iteration (returning to step 2) or finalizes the classifier upon convergence. The same procedure is applied to the HRV dataset, which is split into a training set (19 subjects; 80%) and a validation set (3 subjects; 20%). **<u>Panel C:</u> Retroactive Feature Ablation Analysis:** feature importance is assessed in both datasets to evaluate the impact of individual features on classification performance.

### 2.2 Datasets Transformations (Fig. 1 Panel A)

#### 2.2.1 One Affected Tissue Transcriptome per subject

The vast majority of conventional Transcriptome classifiers typically analyze a single transcriptome derived from the affected tissue of each subject. In order to evaluate the accuracy achievable with traditional classification methods utilizing one sample per individual, we utilized the affected tissue of the datasets and did not utilize the paired control tissue. The BC cohort (21) included 22,279 TMM (trimmed mean of M values) normalized gene expression (22) values from 42 subjects, and the HRV cohort (16) included 20,502 RMA) normalized Affymetrix GeneChip expressions of probe-sets from 19 subjects.

#### 2.2.2 Paired-samples: one Affected tissue transcriptome and one Control tissue per subject

We calculated the fold change by dividing the expression of each mRNA value of the affected tissue by that of the control tissue, in each subject, in each dataset, followed by a log2 transformation (6).

#### 2.2.3 *Single-Subject-Studies (N-of-1-Pathways) are described in* Figures 1 Panel A and Figure 2

### 2.3 Model Selection (Figure 1 Panel A)

We evaluated several machine learning models, including Random Forest (RF), XGBoost, Support Vector Machine (SVM), and Logistic Regression. Random Forest was ultimately chosen due to its robustness, capacity to model non-linear interactions, and superior predictive performance in identifying symptomatic subjects and relevant gene sets (data not shown; available upon request).

**Fig 2.**
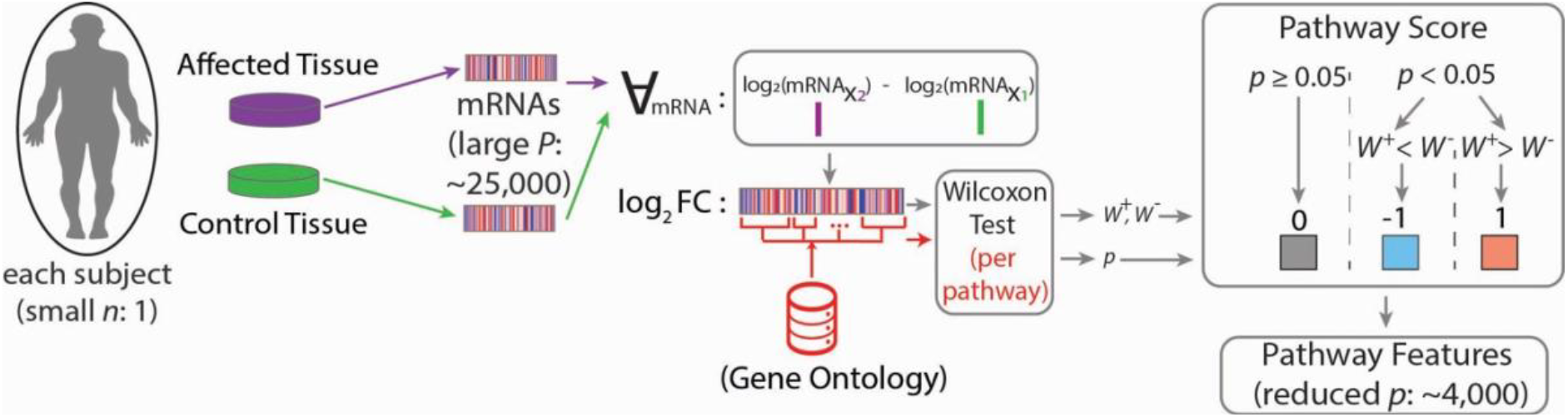
Description of the N-of-1-pathways Wilcoxon analytics in each single subjects. We employed the “N-of-1-pathways” method (17), which aggregates paired RNA-level signals of each subject into pathway-level effect sizes, conducts a non-parametric Wilcoxon test comparing the pathway-associated mRNAs in each Gene Ontology (GO) Biological Processes [pvalue <0.05; other thresholds studied elsewhere (2,3,6,17,16)] for each subject, enabling downstream classification over a smaller number of human-interpretable GO features. This method identifies significantly altered mRNA sets associated to a pathway between two samples of one subject, yielding 4,442 GO mRNA sets in the BC cohort and 2,332 GO mRNA sets in the HRV cohort. The output consists of ternary matrices indicating response status (W-: negatively regulated, W+: positively regulated, and 0: unaltered GO pathway; where W is a significant Wilcoxon test between the one subject’s mRNA levels in the affected tissue sample vs the control among mRNA associated to a specific pathway according to GO; calculated using the ratio Fold Change [FC] of these mRNA expression values). HRV scores were refined with a coefficient of variation <31%.

### 2.4 Classification, Cross Validation, and MLOps (Figure 1 Panel B)

Model robustness was evaluated in both datasets using 5-fold cross-validation. The Random Forest (RF) model was integrated into the Weights & Biases (W&B) MLOps framework (23) to systematically identify features whose interactions significantly contribute to class differentiation. W&B facilitated robust experiment tracking, hyperparameter optimization, and model monitoring, applying consistent hyperparameter ranges across the BC dataset (42 samples) and the HRV dataset (19 samples). This setup allowed us to assess MLOps’ effectiveness in guiding hyperparameter tuning and model tracking while maintaining human oversight. This study was designed to compare the ability of different combinations of data transformations (single-sample per subject, FC, N-of-1-pathways analytics) to improve performance in small human cohorts (small *n* <30 subjects) with high feature dimensionality (very large *p*, transcriptomes = 25,000 mRNA features).

In W&B MLOps, the *sweep.yaml* file configured hyperparameter sweeps by defining key parameters, search strategies, optimization metrics, and other relevant settings for systematic model optimization. Python’s *StratifiedKFold* strategy ensured class proportion consistency across 5 folds and this process was repeated across five iterations with different folds serving as the validation set. Accuracy, precision, and recall performance metrics, were calculated across cross-validation folds and unseen sets (**Tables 2, 3)**. To refine hyperparameter ranges, a human-expert in the loop revised the best hyperparameter intervals using the *sweep.yaml* configuration. This *YAML* file specifies the parameters to be tuned, the search strategy, optimization metrics, and other pertinent settings.

**Table 2.**
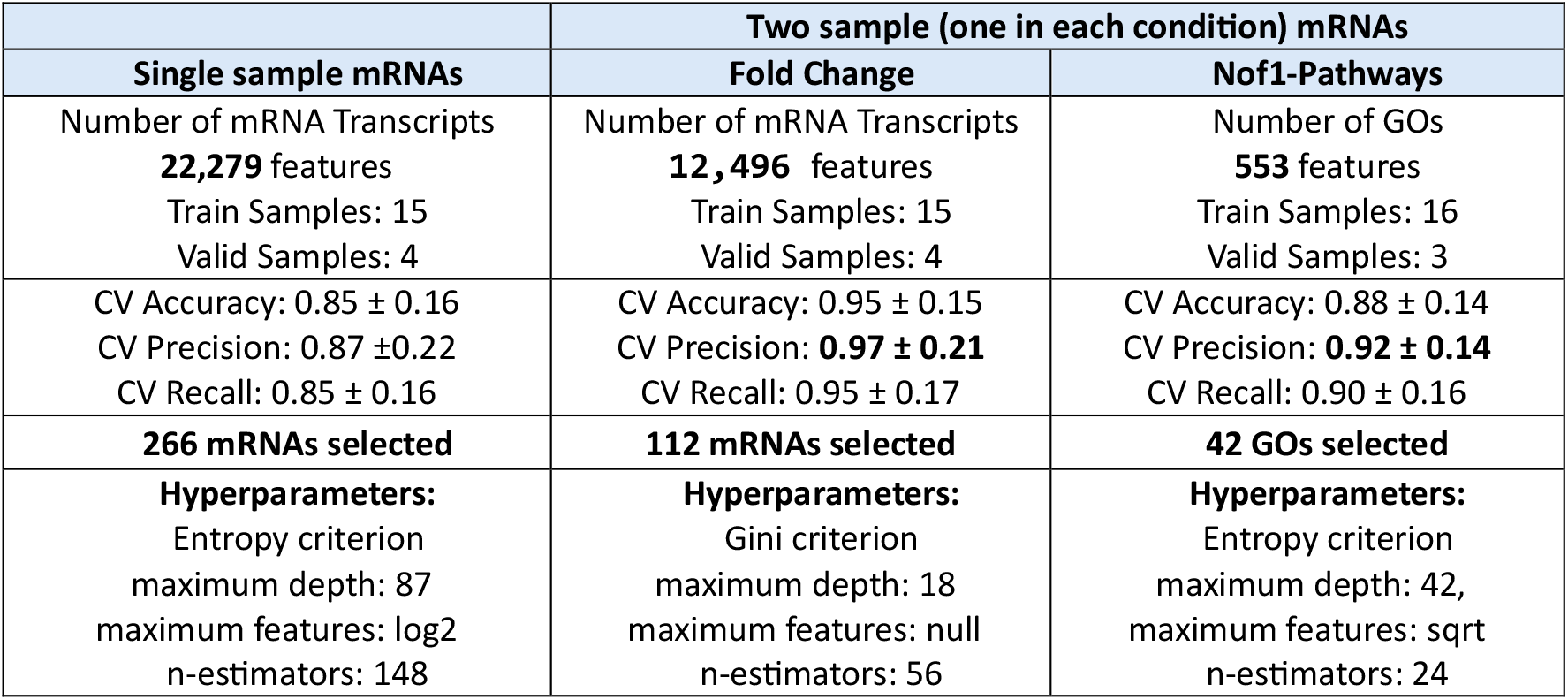
Performance summary of analysis in HRV RF classifier. Legend: CV=cross-validation accuracy ± standard variation, Train=training set, Valid=validation set, Test= test set

**Table 3.**
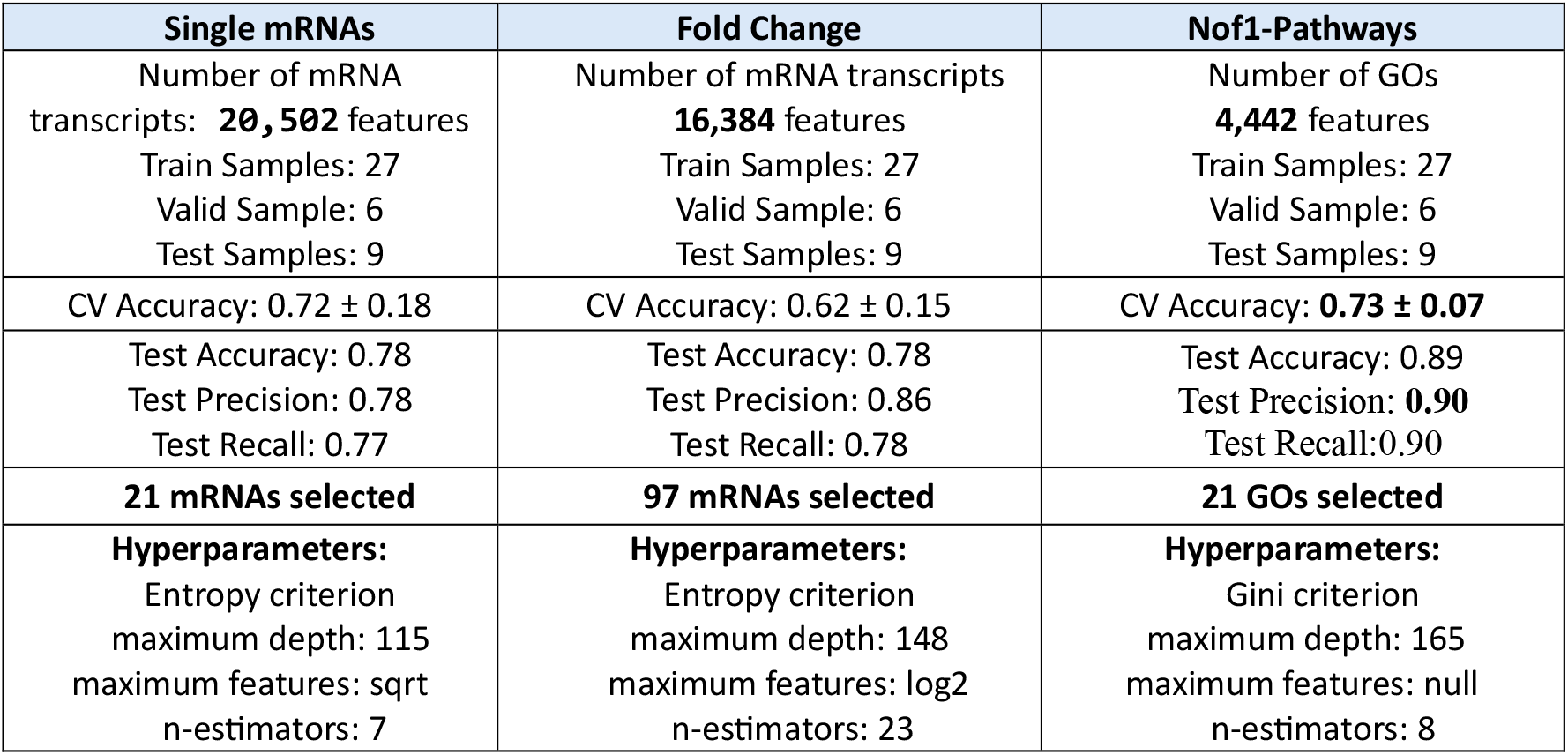
Performance summary analysis in BC RF classifier

| Single mRNAs | Fold Change | Nof1-Pathways |
| --- | --- | --- |
| Number of mRNA transcripts: **20,502** features<br>Train Samples: 27<br>Valid Sample: 6<br>Test Samples: 9 | Number of mRNA transcripts<br>**16,384** features<br>Train Samples: 27<br>Valid Sample: 6<br>Test Samples: 9 | Number of GOs<br>**4,442** features<br>Train Samples: 27<br>Valid Sample: 6<br>Test Samples: 9 |
| CV Accuracy: $0.72 \pm 0.18$ | CV Accuracy: $0.62 \pm 0.15$ | CV Accuracy: **$0.73 \pm 0.07$** |
| Test Accuracy: 0.78<br>Test Precision: 0.78<br>Test Recall: 0.77 | Test Accuracy: 0.78<br>Test Precision: 0.86<br>Test Recall: 0.78 | Test Accuracy: 0.89<br>Test Precision: **0.90**<br>Test Recall: 0.90 |
| **21 mRNAs selected** | **97 mRNAs selected** | **21 GOs selected** |
| **Hyperparameters:**<br>Entropy criterion<br>maximum depth: 115<br>maximum features: sqrt<br>n-estimators: 7 | **Hyperparameters:**<br>Entropy criterion<br>maximum depth: 148<br>maximum features: log2<br>n-estimators: 23 | **Hyperparameters:**<br>Gini criterion<br>maximum depth: 165<br>maximum features: null<br>n-estimators: 8 |

### 2.5 Retroactive Feature Ablation (Figure 1, Panel C)

We *conducted* a feature ablation analysis retroactively on both datasets to assess the impact of the top-ranked features identified by our selected classifiers for each dataset and feature type. This retraining step was conducted to measure the influence of the ablated features on performance metrics such as precision and recall.

## 3 Results (Figure 3, Tables 2 and 3)

In both datasets, RF model robustness was evaluated using 5-fold cross-validation (Methods 2.3-2.4, Figure 1 Panels A-B). For the BC dataset (42 subjects), 80% (27 subjects) were used for training, while the remaining 20% was split into 6 subjects for validation and 9 subjects for testing, ensuring consistent evaluation. Similarly, in the HRV dataset, consisting of 19 subjects, the data was split into 80% (16 subjects) *for* training and 20% (3 subjects) for validation. The *StratifiedKFold* approach from the *scikit-learn* Python package was employed to maintain consistent class proportions across folds, ensuring validation consistency and reproducibility. In MLOps-guided studies (**Methods 2.4, Figure 1 Panel B**), after testing various hyperparameter interval ranges, a human-in-the-loop (expert) confirmed the following optimal RF hyperparameters: criterion (*gini* or *entropy*), number of estimators (5 to 150), maximum features (sqrt, log2, or None), and tree depth (5 to 200). SVM and XGBoost hyperparameters not shown as yielding lower accuracies. Result shown in **Tables 2-3** and **Figure 3**.

**Fig 3.**
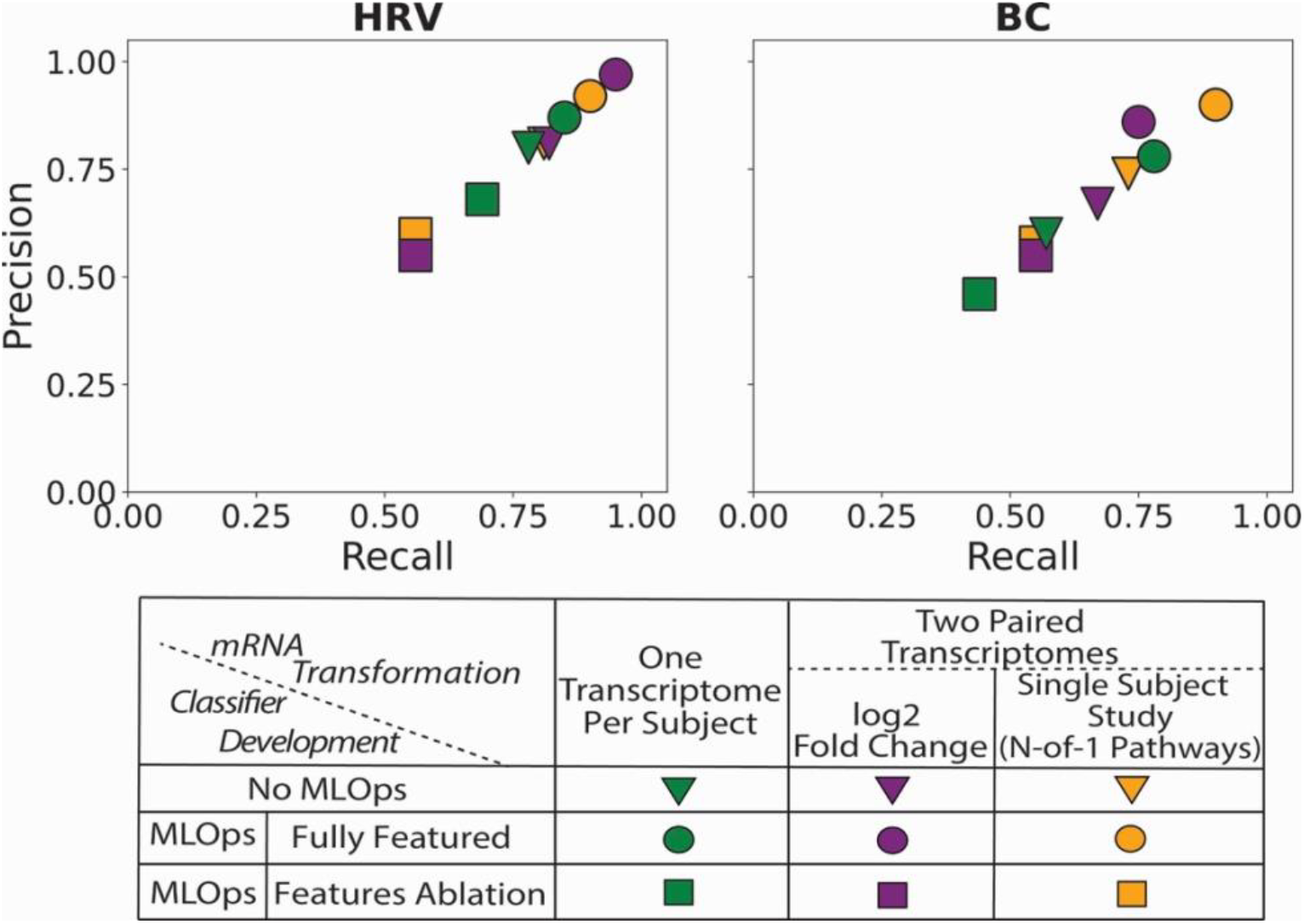
MLOps-guided Precision and Recall according to transcriptome transformation before and after ablation. As shown, paired samples per subject designs (log2FC in **purple**, or N-of-1-pathways single-subject studies in **yellow**) obtained higher accuracies that single sample per subject classification designs (**green**), both in Breast Cancer (BC) and Human Rhinovirus (HRV) micro-cohorts. As expected, retroactive Feature Ablation classifiers (**squares**) were less accurate than the original fully featured classifiers (**circles**). As esxpected, five-fold cross validation classification without MLOps (**triangles**) generated precision and recall that were 14.5% and 12.5% lower.

### Retroactive Feature Ablation Studies in Breast Cancer and HRV Datasets (Figure 3)

To assess the impact of top-ranked features on model performance, an ablation study was performed by sequentially removing the 20 highest-ranked features identified by the classifiers and retraining the optimal Random Forest model with previously tuned hyperparameters. It consisted in masking these features from the data input and re-training (**Methods 2.5**; **Figure 1** Panel C). This analysis quantified the contribution of these features by evaluating changes in precision and recall, revealing a significant decline in predictive accuracy upon their removal. The results underscore the robustness of the selected features derived through the MLOps-driven pipeline, with performance degradation observed across all feature sets.

## 4 Discussion and Limitations

Transcriptome classifiers typically analyze a single transcriptome derived from the affected tissue of each subject. This dataset serves as a baseline to evaluate the accuracy achievable with traditional classification methods utilizing one sample per individual. Specifically, the BC cohort (11) included 22,279 gene expression values normalized using the trimmed mean of M-values (TMM) method (22) from 42 subjects, while the HRV cohort (6) comprised 20,502 Affymetrix GeneChip probe-set expressions normalized using Robust Multi-array Average (RMA) from 19 subjects.

We evaluated several machine learning models, including Random Forest (RF), XGBoost, Support Vector Machine (SVM), and Logistic Regression. Random Forest was ultimately chosen due to its robustness, capacity to model non-linear interactions, and superior predictive performance in identifying symptomatic subjects and relevant gene sets (data not shown).

We performed a retrospective feature ablation analysis on both datasets to evaluate the significance of the top-ranked features identified by our chosen classifiers for each dataset and feature type. This retraining process was crucial to quantify the specific impact of these ablated features on key performance metrics such as precision and recall.

In the HRV micro-cohort, the MLOps-guided fold-change analysis achieved high cross-validated precision (0.97) and recall (0.95). By contrast, single-sample analyses demonstrated a 6% to 8.8% reduction in accuracy and higher susceptibility to noise, primarily due to increased dimensionality. Notably, the MLOps-guided N-of-1 pathways approach exhibited consistent performance, highlighting the efficacy of pathway-level metrics in detecting subtle phenotypic differences within small cohorts. The more heterogeneous BC dataset, characterized by variability in cancer DNA between paired cancer and unaffected tissues, benefited significantly from the N-of-1 pathways analysis, achieving superior test precision (0.90) and recall (0.90) in differentiating TP53 from PIK3CA mutations. Conversely, the MLOps-guided fold-change analysis yielded comparatively lower performance (precision: 0.86, recall: 0.75), suggesting that pathway-anchored features are particularly advantageous for capturing heterogeneous tumor biology.

By systematically applying three mRNA feature transformations (single-sample-per-subject, fold-change, pathways) under identical hyperparameter sweeps within the W&B MLOps platform (*wandb* v0.17.0, Python 3.11.4), our findings emphasize the advantages of paired-sample study designs. However, just as mRNA normalization involves significant engineering expertise, the choice and refinement of feature transformations (fold-change versus pathways) similarly entail expert judgment. The optimization parameters presented (**Tables 2-3**) underscore the synergy between automated methods and expert oversight, highlighting the importance of integrating MLOps with human-in-the-loop analyses for effective micro-cohort classification. **Supplementary Figure 1** suggests these methods enhance biological interpretability and may inform the development of mechanistically meaningful classifiers, although detailed biological analyses are beyond this study’s scope.

Few studies have systematically addressed classifier development requirements in very small cohorts. Our previous work demonstrated feasibility in a prospective cohort (6) without comparative evaluations against conventional methods or MLOps integration. Transfer learning has shown promise in classifying cell types in single-cell RNA sequencing (24) and transcriptomic datasets derived from large human cohorts (25), but these methods have not yet been applied specifically to small human cohorts for clinical predictive analytics.

Several limitations must be noted: (i) alternative machine learning models (SVM, Logistic Regression, XGBoost) consistently underperformed relative to Random Forest, and results were omitted for brevity. Future research should explore fusion deep learning and transfer learning approaches. (ii) Our conclusions are based on limited datasets, necessitating additional transcriptomic data or simulation studies to robustly assess generalizability. (iii) Despite efforts to control overfitting, inherent constraints persist due to small sample sizes, emphasizing the need to develop micro-cohorts through sub-sampling larger paired-sample datasets in future studies; though such datasets are uncommon.

## 5 Conclusion

Most of the approximately 65,000 known human diseases remain inadequately treated due to their rarity and the consequent scarcity of comprehensive studies. The low prevalence of these diseases severely limits conventional transcriptomic approaches, as bulk RNA sequencing (bRNAseq) typically necessitates larger cohorts for effective classifier development. Emerging technologies such as spatial RNA sequencing and single-cell RNA sequencing present promising alternatives suitable for smaller cohort studies; however, these methods currently incur approximately 20 times higher costs per sample and capture around five times fewer mRNA transcripts. As these technologies become more affordable and achieve improved transcriptomic coverage, novel analytical methodologies tailored for small cohorts are expected to evolve. Additionally, transfer learning techniques, already successfully applied to large-scale transcriptomic datasets, offer considerable potential for small-cohort classification. However, standardized frameworks for applying transfer learning specifically to paired-sample designs are not yet established, highlighting an important area for future research.

This study systematically evaluated multiple complementary approaches designed to enhance the statistical power of bulk mRNA-based classification within micro-cohorts. Our results demonstrate that the integration of these approaches improves precision and recall by approximately 12.5%–14.5% compared to traditional single-sample methodologies. Specifically, we propose strategies that include: (i) leveraging paired comparisons of affected and control tissues within individual subjects, and (ii) employing MLOps-guided analytical workflows combined with expert-in-the-loop oversight to ensure robustness, transparency, and reproducibility. This paired-sample methodology has been shown to improve classifier development in large cohorts (6, 26-31), and here we show that, in very small cohorts, it also facilitated classifier development at both the individual mRNA level (via fold-change analysis) and the biologically interpretable knowledge-anchored pathway level (through N-of-1 pathway-based analyses leveraging Gene Ontology gene sets). The adoption of MLOps practices optimized hyperparameter tuning and model deployment, while expert oversight ensured the biological validity of the results. Collectively, these strategies effectively address the challenges posed by high feature dimensionality and limited sample sizes, thereby laying the groundwork for advancing personalized therapeutic interventions in rare disease contexts.

Although this investigation primarily targeted a specific biological scale, optimal classifiers for clinical prediction are likely to incorporate comprehensive real-world data across diverse biological and clinical dimensions. Future research endeavors should integrate transcriptomic data with multi-omics approaches, medical imaging, and patient-centric outcomes to further enhance predictive accuracy and personalized medicine capabilities.

## Acknowledgements

All authors contributed to the analysis and interpretation of the results. Study design was performed by MS, NP, and YAL. Machine learning analyses were conducted by MS. N-of-1 pathways transformation was completed by LSW. Figure contributions: Figure 1 by MS; Figure 2 by MS, LSW, NP, and YAL; Figure 3 by MS, LSW, and YAL. Table contributions: Table 1 by NP and YAL; Tables 2 and 3 by MS and YAL. This study was partially funded the University of Utah.

## Conflict of interest

The authors have no conflict of interest.

## Disclosures

All authors have seen and approved the manuscript, and that it hasn’t been accepted or published elsewhere in its entirety (a poster proceeding is disclosed below). The authors declare no conflicts of interest. Large language models (LLMs) were employed **solely** for grammatical editing and improving manuscript clarity. Portions of the breast cancer results (a subset of Table 3 and of the BC panel of Figure 3) were previously presented as a poster titled “Pathway-Anchored Dimension Reduction and MLOps Enable Robust Micro-Cohort Genome-by-Environment Classifier Development” at the American Medical Informatics Association (AMIA) Informatics Summit, March 9–13, 2025, Pittsburgh, PA, USA. Additionally, only the subset of finding pertaining to Table 3 and the subpanel “ BC” of Figure 3 were published in the proceedings of the 23rd International Conference on Artificial Intelligence in Medicine (AIME 2025), June 23–26, 2025, Pavia, Italy: Shabanian M, Pouladi N, Wilson LS, Prosperi M, Lussier YA. Enabling Transcriptome Classification in Micro-Cohorts with Pathway-anchoring and Single-Subject Studies, Springer, Artificial Intelligence in Medicine Proceedings, Lecture Notes in Computer Sciences Series, Vol.15735 (in press).

---

**Suppl. Fig. 1.**
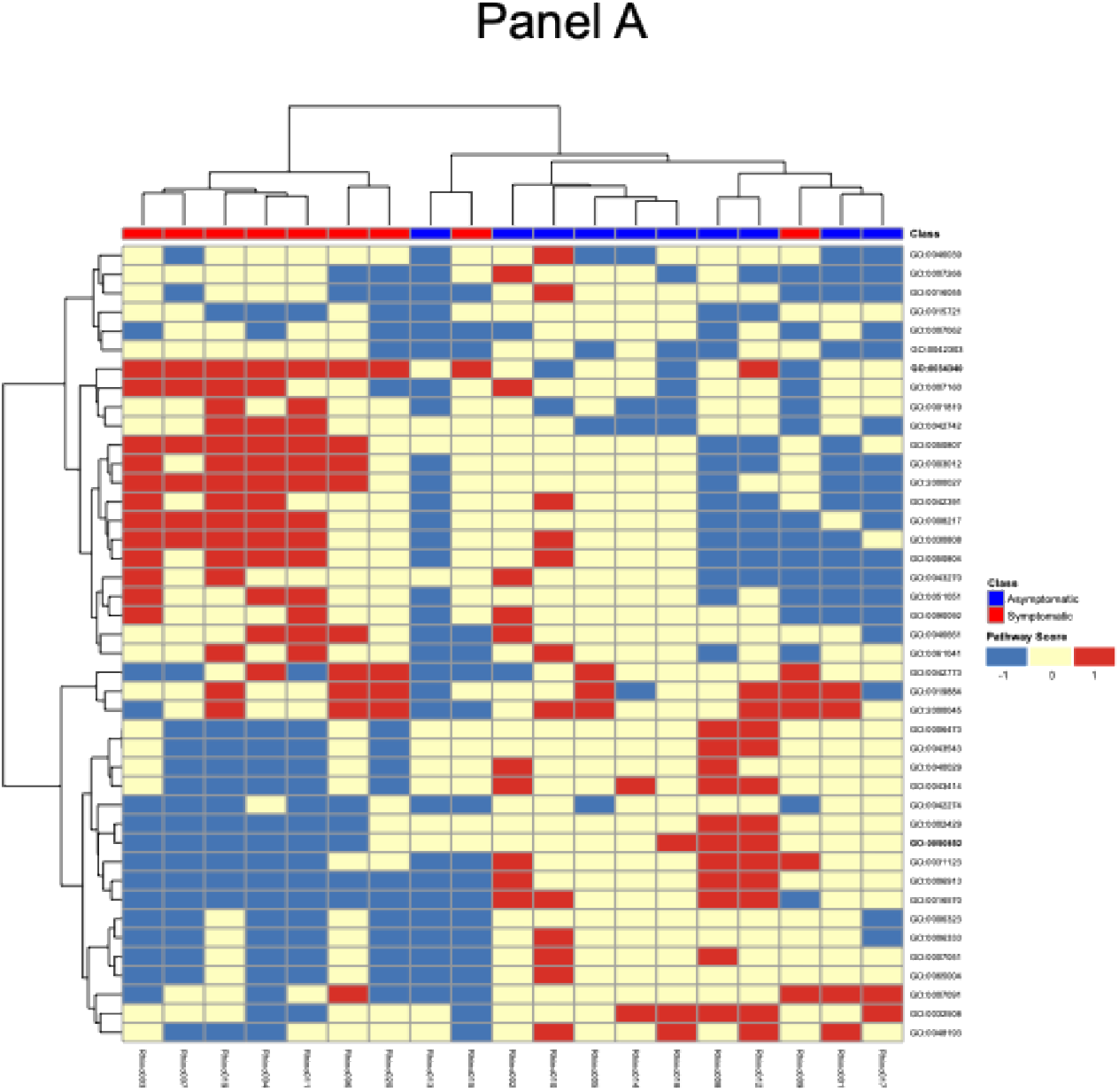
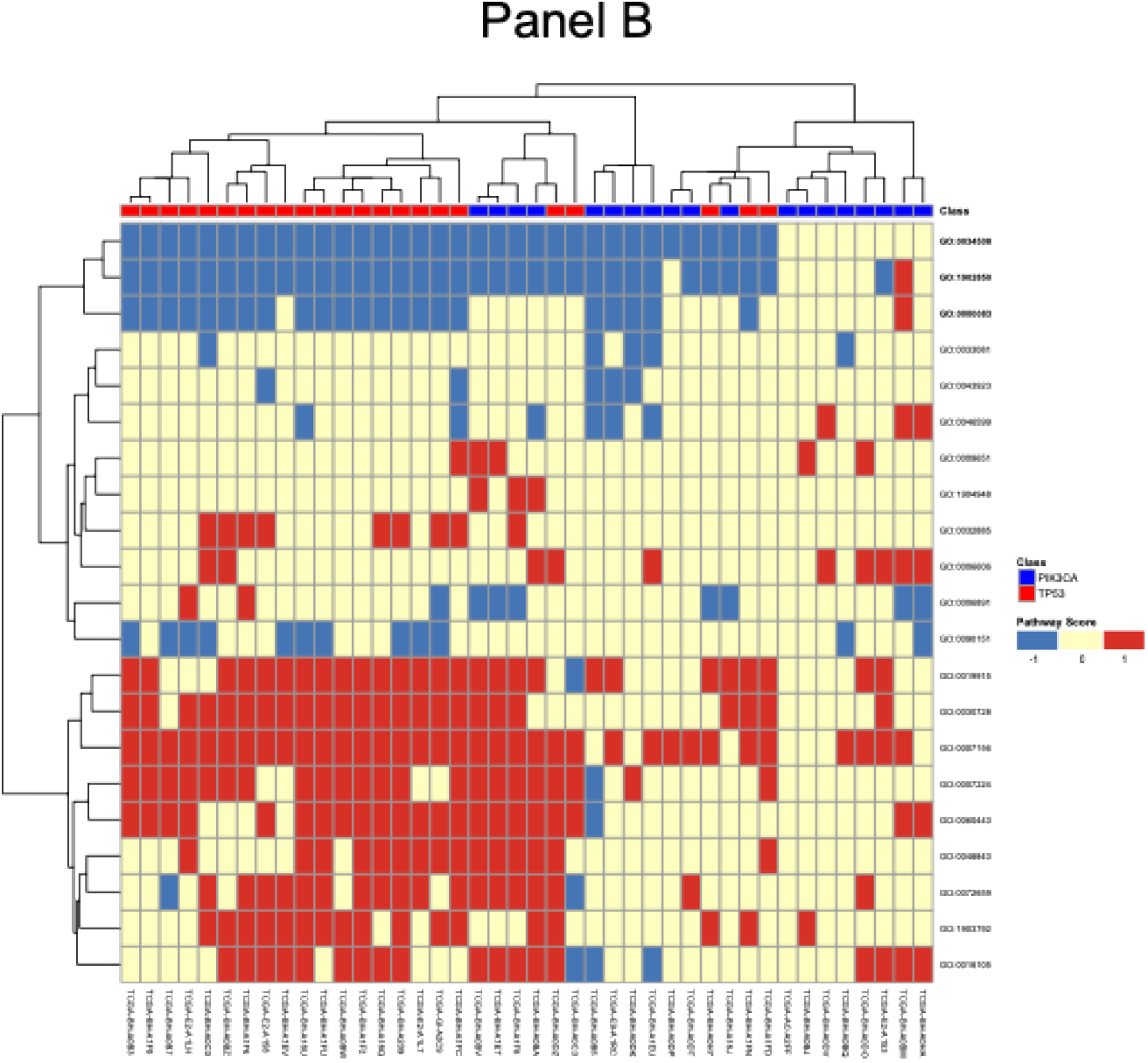
MLOps-anchored pathways-level studies suggest increased human interpretability over mRNA transcript ones. MLOps-guided pathway-level analyses offer improved biological interpretability over conventional mRNA-level approaches. Panel A highlights immune response divergence in HRV infections, while Panel B reveals distinct cell cycle regulation patterns between PIK3CA- and TP53-driven breast cancers, aligning with known oncogenic mechanisms. Heatmaps were generated using the R package Pretty Heatmaps with Ward.D2 clustering to visualize patient segregation into phenotypic groups based on N-of-1 scores of the prioritized GO biological processes (see Figure 2). ***Panel A. HRV symptomatic vs asymptomatic infections***. Among the top 4 biological processes by classifier importance, two show that the ***innate immune response*** (*GO:0034340 interferon response*) is significantly increased in symptomatic subjects (W^+^ for 8 out of 9) and either negative (W^-^ for 2 out of 10) or unchanged in asymptomatic patients when comparing their pathway mRNAs levels during HRV infection and before (Wilcoxon test in each pathway of each subject). We observe the reverse for the ***adaptive immune response*** (*GO:0050852 T cell receptor signaling*) significantly downregulated in 6 out of 9 symptomatic subjects (W ^-^) and Wilcoxon test not significant in the remainder, while and either positive (W^+^ for 3 out of 10) or unchanged in asymptomatic patients **Panel B. *PK3CA-vs TP53-“oncogene-driven” Breast Cancers***. PIK3CA and TP53 are very common across various subtypes of invasive breast cancer, thus discriminating their downstream effects are of utmost clinical importance to narrow down the breast tumor heterogeneity. Our findings show that more than 90% patients with TP53 mutation as opposed to 45% of those with PIK3CA mutation show lower activity in the top three prioritized gene sets by Random Forest (*GO:0000083: regulation of transcription involved in G1/S transition of mitotic cell cycle, GO:0034508: centromere complex assembly and GO:1902850: microtubule cytoskeleton organization involved in mitosis*) that underpin biology of the ***cell cycle*** and thus ***progression of tumor***. This is consistent with biological downstream effect of TP53 as a tumor suppressor.

## Notes

### Competing Interest Statement

The authors have declared no competing interest.

